# Combining gene network, metabolic, and leaf-level models show means to future-proof soybean photosynthesis under rising CO_2_

**DOI:** 10.1101/582981

**Authors:** Kavya Kannan, Yu Wang, Meagan Lang, Ghana S. Challa, Stephen P. Long, Amy Marshall-Colon

## Abstract

Global population increase coupled with rising urbanization underlies the predicted need for 60% more food by 2050, but produced on the same amount of land as today. Improving photosynthetic efficiency is a largely untapped approach to addressing this problem. Here, we scale modeling processes from gene expression through photosynthetic metabolism to predict leaf physiology in evaluating acclimation of photosynthesis to rising [CO_2_]. Model integration with the yggdrasil interface enabled asynchronous message passing between models. The multiscale model of soybean photosynthesis calibrated to physiological measures at ambient [CO_2_] successfully predicted the acclimatory changes in the photosynthetic apparatus that were observed at 550 ppm [CO_2_] in the field. We hypothesized that genetic alteration is necessary to achieve optimal photosynthetic efficiency under global change. Flux control analysis in the metabolic system under elevated [CO_2_] identified enzymes requiring the greatest change to adapt optimally to the new conditions. This predicted that Rubisco was less limiting under elevated [CO_2_] and should be down-regulated allowing re-allocation of resource to enzymes controlling the rate of regeneration of ribulose-1:5 bisphosphate (RubP). By linking the GRN through protein concentration to the metabolic model it was possible to identify transcription factors (TF) that matched the up- and down-regulation of genes needed to improve photosynthesis. Most striking was TF GmGATA2, which down-regulated genes for Rubisco synthesis while up-regulating key genes controlling RubP regeneration and starch synthesis. The changes predicted for this TF most closely matched the physiological ideotype that the modeling predicted as optimal for the future elevated [CO_2_] world.

## INTRODUCTION

As the world’s most important seed legume and most widely grown dicotyledonous crop, the future-proofing of photosynthesis in soybean (*Glycine max* (L.) Merr.) under rising atmospheric concentrations of CO_2_ ([CO_2_]) is of importance. Down-regulation of light-saturated net leaf CO_2_ uptake (*A*_*sat*_) at elevated [CO_2_] has been reported for many C_3_ crops, yet the mechanism underlying this response is poorly understood. Under current [CO_2_], *A*_*sat*_ in C_3_ crops is most commonly limited by the *in vivo* Rubsico activity (*V*_c,max_) (Long et al., 2004). However, as [CO_2_] continues to rise, it follows from the steady-state biochemical model of photosynthesis of (Farquhar et al., 1980) and its subsequent modifications (Von Caemmerer, 2000) that control will shift from Rubisco to RubP regeneration (Long et al., 2004), which is represented by the maximum *in vivo* rate of whole chain electron transport (*J*_max_). While described by electron transport, most evidence now points to this being limited by the metabolic steps of carbon metabolism leading to RubP regeneration (Raines, 2003, Stitt and Sonnewald, 1995). This shift from Rubisco-to RubP-limited photosynthesis permits a reduction in leaf Rubisco content without a loss in *A*_sat_ (Woodrow, 1994, Long et al., 2004, Ainsworth and Long, 2005). Because Rubisco accounts for the largest single share of leaf N, optimization of Rubisco content would maximize the efficiency of use of this commonly limiting resource (Drake et al., 1997, Long et al., 2004).

When [CO_2_] is increased around a photosynthesizing leaf, *A*_*sat*_ can increase for two reasons, first the *K*_M_ of Rubisco for CO_2_ is close to the current atmospheric concentration, so elevated [CO_2_] increases the velocity of carboxylation, and secondly, CO_2_ competitively inhibits the oxygenation reaction that produces phospho-glycolate and in turn photorespiration. This latter effect is particularly important because it increases the efficiency of net CO_2_ uptake by diverting ATP and NADPH (generated by the light reactions) away from photorespiratory metabolism to photosynthetic assimilation, and so will increase *A*_sat_ regardless of other limiting factors. Under rising [CO_2_] both factors will increase *A*_sat_ when *V*_c,max_ is limiting, but only the second factor when *J*_max_ is limiting. Assuming the average specificity and *K*_M_ for CO_2_ and O_2_ for Rubisco from terrestrial plants, and a constant intercellular versus external [CO_2_], one can calculate the increase in *A*_sat_ that would result from an increase in atmospheric [CO_2_]. Following the procedure of (Long et al., 2004) for a leaf temperature of 25 °C, the increase in atmospheric [CO_2_] from today’s 400 µmol mol^−1^ to 550 µmol mol^−1^ would increase Rubisco-limited and RubP-limited photosynthesis by 31% and 9%, respectively. 550 µmol mol^−1^ is the concentration forecast for 2050 assuming current emissions trends continue (RCP8.5, (Pachauri et al., 2014)). At current atmospheric [CO_2_] soybean leaf photosynthesis is at the transition point between *V*_c,max_- and *J*_max_-limitation (Bernacchi et al., 2005). Therefore, as [CO_2_] rises soybean photosynthesis would become RubP-limited. If however, resource currently invested in Rubisco was re-allocated to increased *J*_max_ then this transition point would move to a higher [CO_2_] and a 31% rather than 9% increase in *A*_sat_ could be obtained, without any increased total investment of protein in the photosynthetic apparatus. When grown under elevated [CO_2_] in open-air field conditions, is an increase in J_max_ observed at the expense of V_c,max_?

In two consecutive years, (Bernacchi et al., 2005) analyzed photosynthesis in a modern highly productive soybean cultivar under open-air [CO_2_] elevation using Free-Air Concentration Enrichment (FACE) (Long et al., 2006). Compared to control plants those grown in [CO_2_] elevated to 550 µmol mol^−1^ showed a shift in control of A_sat_ from co-limitation of V_c,max_ and J_max_ to limitation solely by RubP-limitation. There was a significant 5% decrease in the ratio of V_c,max_:J_max_, showing a decline in Rubisco activity relative to the capacity for RubP regeneration (Bernacchi et al., 2005). However, while acclimation had occurred it was insufficient to maximize the potential increase in A_sat_, had the system responded to fully re-optimize investment of resources. At 25 °C, re-optimizing the system to 550 ppm was calculated to require a 35% reduction of investment in Rubisco and re-allocation of that protein to the apparatus for regeneration of RubP (Drake et al., 1997, Long et al., 2004), while only a 5% change was observed. The plant was apparently over-investing in Rubisco and under-investing in the apparatus determining regeneration of RubP. How might genetic manipulation be used to achieve re-optimization and prepare soybean and other crops to sustainably maximize photosynthetic efficiency and in turn crop productivity under future conditions?

Here we combine a metabolic model of C_3_ photosynthetic metabolism, including the C_2_ photorespiratory pathway, mathematically representing all discrete steps of photosynthesis from light and CO_2_ absorption to starch and sucrose synthesis (Zhu et al., 2007, Zhu et al., 2013) with a gene network model to predict observed acclimatory changes. This is successfully tested against observed acclimatory changes of photosynthesis in soybean grown at elevated [CO_2_]. Finally, via sensitivity analysis and dynamic gene regulatory network analysis, the combined model is used to predict genetic changes, including expression levels of transcription factors, that could fully optimize leaf photosynthetic efficiency to future elevated [CO_2_] conditions.

## MULTISCALE MODEL DEVELOPMENT

To predict what genes may transcriptionally regulate the soybean response to elevated [CO_2_], it was necessary to develop a mechanistically informed model in which the multi-scale response could be explored. We have previously developed complete mechanistic metabolic models of photosynthetic carbon metabolism that successfully predict dynamic responses of leaf chlorophyll fluorescence and fluxes of CO_2_ and O_2_ to changes in light, [CO_2_] and [O_2_] (Zhu et al., 2007, Zhu et al., 2013). While each of these models provided new insights about photosynthesis, when combined with optimization routines to predict optimal investments for different environments, they are not equipped to predict transcriptomic and genetic changes that could achieve those optimal patterns of investment. The generalization of whole plant metabolism and signaling pathways often results in simulations with low prediction accuracy upon model perturbation. Multiscale models that mimic the biological flow of information across scales have been shown to have higher prediction accuracy than models at individual scales, especially when simulating conditions different from the original training data (Chew et al., 2014). To our knowledge, no current model of photosynthesis in soybean scales from genes to organs. Such a model could potentially simulate system-wide changes in photosynthesis in response to targeted genetic perturbations.

To predict leaf-level responses of net CO_2_ uptake, a metabolic model (e-Photosynthesis) was combined with a leaf micrometeorological model that integrated boundary layer conductance, stomatal conductance, and leaf energy balance (Drewry et al., 2010, Nikolov et al., 1995). Our prior e-Photosynthesis model (Zhu et al., 2013) simulates fluxes through some 70 reactions involved in the light and dark reactions of C_3_ photosynthesis. The steady-state photosynthesis rate predicted by the e-Photosynthesis model replaced the leaf model prediction that was based on the Farquhar model of photosynthesis (Nikolov et al., 1995). Leaf temperature, light intensity and intercellular CO_2_ concentration predicted by the leaf model were used as inputs for the e-Photosynthesis model. The integrated leaf metabolic and micrometeorological model effectively simulates the observed response to net leaf CO_2_ uptake to [CO_2_] observed for soybean in the field (Bernacchi et al., 2005) (Figure 1, Supplemental figure 1). However, it lacks any capacity to predict the observed decrease in *V*_c,max_ and increase in *J*_max_ that resulted from growth in elevated [CO_2_]. This is because the model lacks any connection to the underlying changes in gene expression that may cause this altered photosynthetic phenotype.

**Figure 1.**
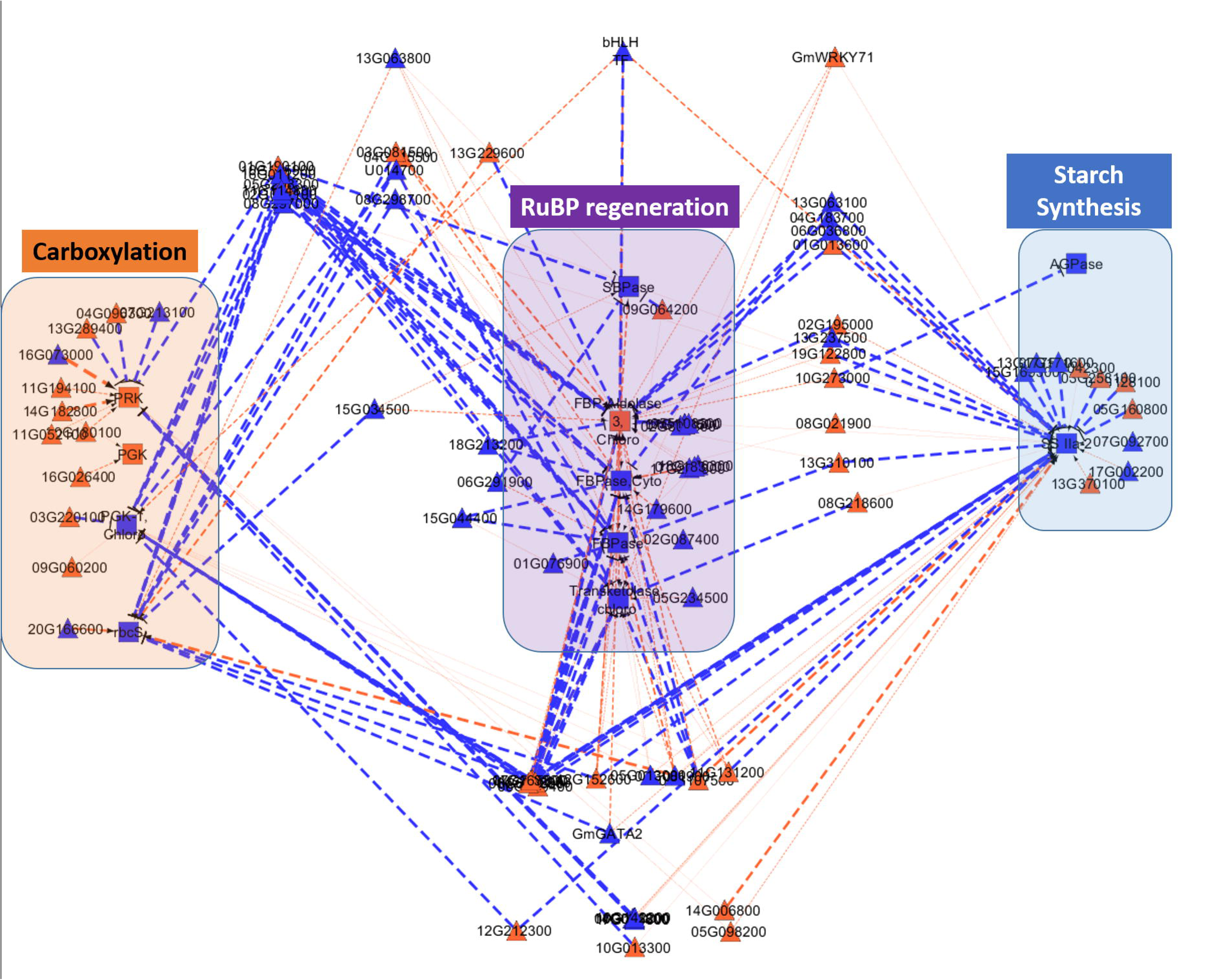
Simulated and measured (Bernacchi et al., 2005) photosynthetic carbon dioxide response curves of soybean growing in ambient CO_2_ (370 μmol mol^−1^) and elevated CO_2_ (550 μmol mol^−1^). PAR is 2000 µmol m^−2^ s^−1^

The altered photosynthetic phenotype likely resulted from adjusted enzyme concentrations in soybean leaves grown under ambient and elevated CO_2_ concentrations. Such CO_2_-induced changes in protein concentrations was shown for total protein concentration in barley, rice, wheat, soybean, and potato (Taub et al., 2007) and for specific proteins via proteomic analysis in rice (Bokhari et al., 2007), wheat (Yousuf et al., 2017), and grape (Zhao et al., 2019). A decrease in the quantity of Rubisco is a pervading feature of plants grown in the field under elevated [CO_2_] (Ainsworth and Long, 2005). However, the e-Photosynthesis model only allows substrate (CO_2_) concentration to change, which results in altered reaction rates, but lacks capacity to predict acclimatory changes in enzyme concentrations. By including gene expression data from soybean plants grown under ambient and elevated [CO_2_] (Leakey et al., 2009), it is possible to make predictions about changes in enzyme concentrations.

Gene expression data cannot be used as direct input for the e-Photosynthesis model, which can only accept enzyme concentrations. Also, transcriptome data from microarray or RNA-sequencing technologies provide relative and not absolute quantification of mRNA transcripts. To overcome these challenges and inform the e-Photosynthesis model with gene expression data, a model was needed to computationally translate mRNA to protein concentration. An ordinary differential equation (ODE) was adapted from (Becskei and Serrano, 2000), and given as:

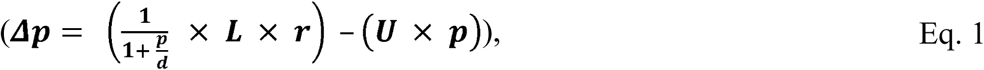

where, *L* and *U* are the estimated gene family-specific protein synthesis and degradation rates (Li et al., 2017), respectively, *r* is the mRNA level, *p* is the initial protein concentration, and *d* is the upper limit of protein translation. It is assumed that *r* and *p* are equal (Edfors et al., 2016), and *p* is based on starting protein concentration estimates from the e-Photosynthesis model. To simulate steady-state protein concentrations in elevated CO_2_, *p* was adjusted based on the proportion of change in mRNA level between ambient and elevated [CO_2_] for a given gene.

The change in predicted, relative protein concentrations between ambient and elevated CO_2_ conditions (ProteinRatio) was used to adjust the enzyme concentration and activity of each gene involved in the light and dark reactions of photosynthesis represented in the e-Photosynthesis model, as follows:

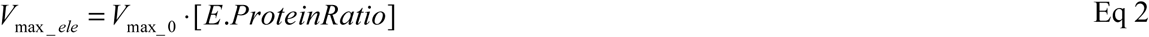

where, *V*_*max_ele*_ is the maximum activity of each enzyme in elevated CO_2_, *V*_*max_0*_ is the original maximum activity of each enzyme in ambient CO_2_, and *E* is the estimated enzyme concentration.

Because the protein translation model and metabolic/leaf level models are implemented in different programing languages (Python and MATLAB) respectively, they cannot communicate with each other directly without significant alteration of the model code or through manual integration by running the model programs independently and using files to pass information between them. In order to integrate the models programmatically (Figure 2), we used the yggdrasil framework. yggdrasil is an open-source Python package developed by the Crops *in silico* research group for connecting models written in different programming languages through simple interfaces in the model’s language of implementation. Based on information on the target models and connections between the models provided in human-readable specification files, yggdrasil runs the designated models in parallel and coordinates asynchronous communication between the models as they run. Asynchronous message passing allows models to continue working after sending output to the next model in the network without waiting for the output to be received, thereby improving the performance of the overall model integration as models can complete calculations simultaneously in separate processes. yggdrasil currently supports connecting models written in Python, Matlab, C and C++ on Linux, Mac OS, and Windows operating systems. Additional information on the yggdrasil package can be found in (Lang, 2019) or in the documentation (https://cropsinsilico.github.io/yggdrasil/).

**Figure 2.**
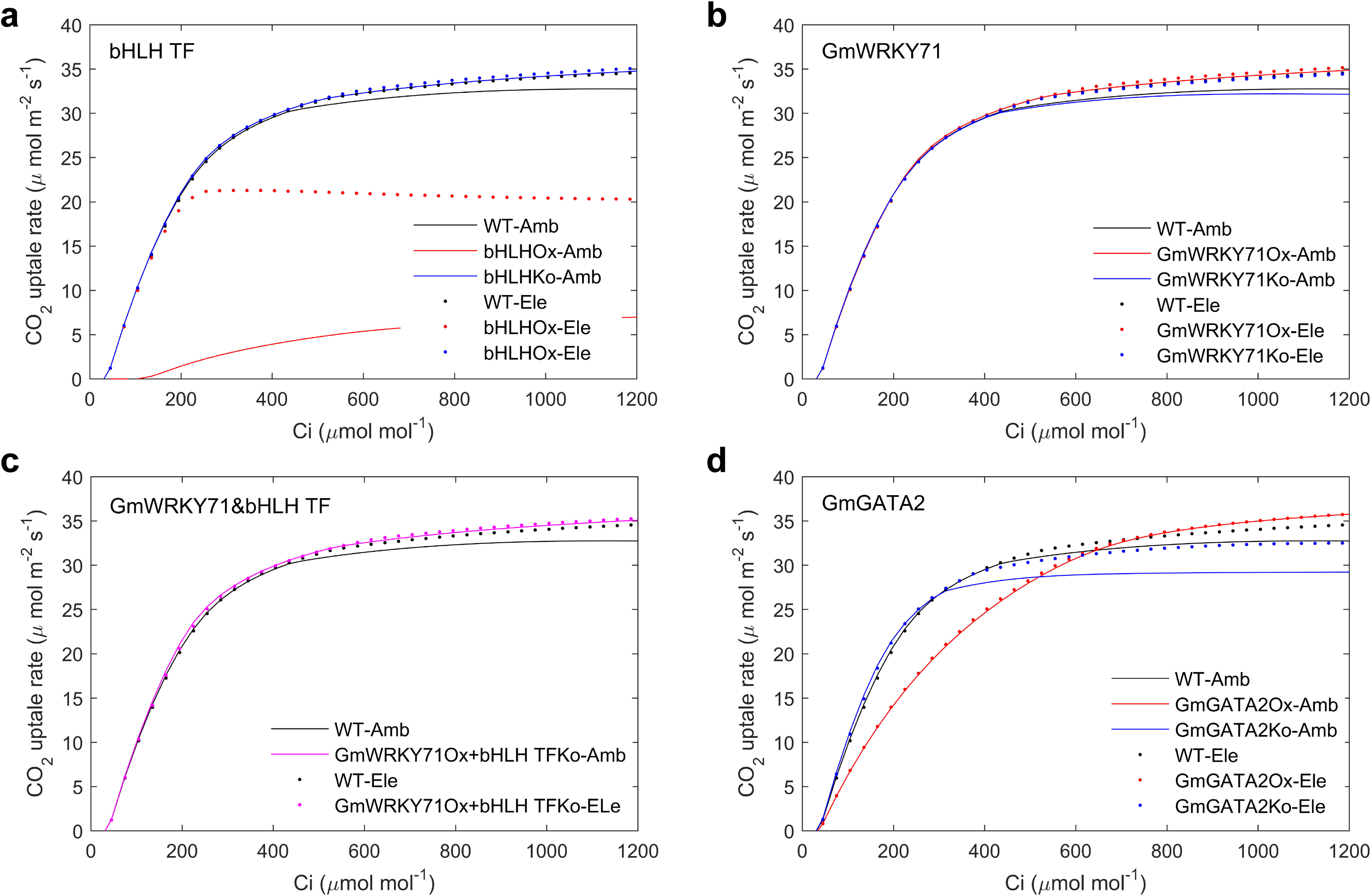
Model integration schematic describing the scaling from the gene regulatory network model to the metabolic model to the leaf physiological model. Note that these models interact through state variable indicated in the arrows. Ca is ambient CO_2_ concentration, PPFD is the amount of light absorbed by the leaf, T is leaf temperature and A is the net carbon assimilation.

## METHODS

### Differential Expression Analysis of Genes Responding to CO_2_

The soybean transcriptome differential responses to growth in ambient and elevated [CO_2_] was obtained previously using the Affymetrix *Glycine max* genechip (Leakey et al., 2009). These data were re-analyzed here to identify differentially expressed (DE) genes corresponding to leaf tissue collected at the beginning of seed set when the canopy has attained full maturity. Probe sets were normalized using the GC Robust Multi-array Average (GCRMA) method. A one-way ANOVA was done in the R statistical software environment using the aov function of the stats package. The reanalysis identified 5005 DEGs on a statistical cutoff of P value <= 0.05 and Benjamini-Hochberg False Discover Rate (FDR) <= 0.3.

### Construction of CO_2_ responsive Gene Regulatory Network

#### Co-expression using mutual rank

Mutual rank analysis (Obayashi and Kinoshita, 2009) was used to obtain highly significant co-expression relationships among differentially expressed genes (DEGs). In mutual rank analysis, the Pearson Correlation Coefficient (PCC) is calculated between the gene of interest and all other DEGs, then sorted based on their PCC ranks, in which the gene pair having the highest correlation value is given rank 1 (Obayashi and Kinoshita, 2009). Mutual Rank is calculated from PCC rank by taking the geometric mean between PCC ranks from gene A to gene B and from gene B to gene A, as these can be different. The ranks are scaled between 0 to 1, where MR of 1 indicates the most significant coexpression interaction. All interactions having an MR >= |0.8| and a PCC >= |0.6| were selected for the coexpression network. Interactions having a significant correlation were further filtered to retain only those that had predicted binding interactions between the TF and target gene as described below.

#### Static Gene Regulatory Network construction and analysis

To analyze gene regulatory networks, a DNA pattern search algorithm was performed (PlantPAN, http://plantpan2.itps.ncku.edu.tw/) (Medina-Rivera et al., 2015) to identify the presence of known *Cis*-regulatory elements (CREs) in the promoter region of genes of interest. Known CREs were obtained from the transcription factor databases Transfac (Matys et al., 2006), JASPAR (Sandelin et al., 2004), CISBP (Weirauch et al., 2014), and PlantTFDB (Jin et al., 2013), and NewPLACE (Higo et al., 1999). The promoter region considered in this study was the sequence 1 Kb upstream of the predicted or experimentally verified Transcription Start Site (TSS) for every gene obtained from PlantProm (Shahmuradov et al., 2003). Promoter regions of target genes were analyzed for an enrichment of CREs for a particular TF family. For this analysis, the average number of binding sites for a CRE in the putative 1 Kb up-stream promoter region of all the genes present in Soybean Wm82.a1.v1 was calculated. If a target gene promoter has a greater than average number of TF family-specific binding sites present in its promoter, then the CRE of interest is considered over-represented.

### Dynamic Gene Regulatory Network Model Construction

A dynamic Gene Regulatory Network (GRN) model was built using a linear regression algorithm to infer relationships between a dependent variable (in this case the expression of the putative target gene) and one or more independent variables (or predictors; in this case TFs). The resulting linear model was used to predict the response variable based on the states of the dependent variables. The regression algorithm was run in R using the LM function (Team, 2013), which optimizes variables of the linear model using a least squares fit between the response and dependent variables on training data (Eq 1) (Bjorck, 1996).

For example:

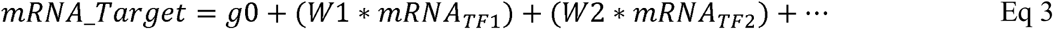

where, g0, W1, W2 are least squares optimized parameters for the linear model. mRNA_TF1, mRNA_TF2, etc. are expression values of TFs predicted to regulated target genes of interest in the static GRN. Parameters were optimized using training data, which ultimately resulted in a weight (W_x_) that corresponds to the level of influence that a TF exerts on a predicted target gene’s expression. A linear model was generated for every gene in the static CO_2_-responsive GRN, and used to simulate the expression of genes of interest in both ambient and elevated CO_2_ environments. All linear model equations are listed in Supplemental Table 1.

The training data was obtained from seven soybean Affymetrix microarray transcriptome datasets (GSE8432, GSE23129, GSE26198, GSE29740, GSE29741, GSE35427, GSE44685). While these datasets were derived from a variety of experimental conditions, they were chosen because samples were taken from similar tissue and developmental stage as those in the CO_2_- responsive dataset used to build the static GRN (Leakey et al., 2009). Expression data from all training sets were commonly normalized by GCRMA and quality control analysis was performed; samples with a high variation in their median expression level within replicates were removed from the analysis. A total of 213 samples were used to train the linear regression model. The CO_2_-responsive dataset that is used to build the static GRN was used as a test dataset for the linear model, to predict expression of the target gene using the optimized weight associated with every TF.

### Protein translation model

A protein translation model (PTM) adapted from (Becskei and Serrano, 2000) was used to predict steady-state protein concentrations based on relative mRNA transcript levels (See Eq. 1) The model parameter *p* (and thus *r*) is obtained from the initial protein concentration used in the e-Photosynthesis metabolic model under the control (ambient CO_2_ = 380 ppm) condition. To predict initial protein concentration in elevated CO_2_ conditions (550 ppm), *p* was adjusted using the relative fold change in mRNA transcript levels measured between ambient and elevated CO_2_ conditions. As stated earlier, *L* and *U* are protein synthesis and degradation rates, respectively, and denotes the increase or decrease in protein abundance per hour (denoted as g/L/hour). Soybean gene-specific *L* and *U* rates were estimated based on the rates of their Arabidopsis orthologs taken from (Li et al., 2017). If ortholog information for a gene was not available, *L* and *U* were estimated by taking an average of *L* and *U* rates for all Arabidopsis genes involved in photosynthesis.

The PTM model simulations resulted in steady-state protein concentration ratios between ambient and elevated [CO_2_] conditions for every gene. Optimized parameter ‘*d*’ was obtained in a gene specific manner such that, the steady-state protein concentration ratio between the two conditions (elevated/ambient) remains constant for that gene after a particular threshold ‘*d*’ (see Supplemental Table 2). This protein concentration ratio was then used as one of the inputs for the e-Photosynthesis model (described below) in order to account for changes in gene expression as a factor influencing the enzyme kinetics of proteins in the primary C metabolism machinery. The model assumes constant temperature and constant concentration of RNA polymerase (El Samad et al., 2005).

### Metabolic photosynthesis model

The soybean photosynthesis metabolic flux (MF) model is based on the e-Photosynthesis model (Zhu et al., 2013) and implemented in MATLAB. The e-Photosynthesis model is a general C_3_ photosynthesis model that includes each discrete process from light capture to carbohydrate synthesis, including photorespiratory C_2_ metabolism. In the model, the rate of change of the concentration of each metabolite is represented by an ordinary differential equation (ODE):

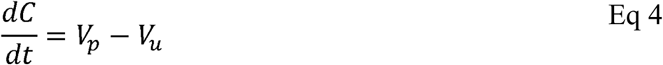

Where, [C] represents metabolite concentration; *V*_*p*_ is the total rate of reaction(s) that produces C, and *V*_*u*_ is the rate of consumption. Rate equations for each enzyme catalyzed reaction were developed based on standard Michaelis-Menten equations for enzyme kinetics, with kinetic parameters corresponding to a temperature of 25°C. Four enzymes, including Rubisco, did not satisfy the conditions needed to apply Michaelis-Menten kinetics, and equations for their catalysis were as in (Zhu et al., 2007). Initial protein concentration of enzymes in the MM kinetics equation were obtained from (Zhu et al., 2013) (Supplemental Table 3). *V*_c,max_ and *J*_max_ of soybean grown under ambient and elevated [CO_2_] in the field had been determined previously (Bernacchi et al., 2005). This was the same germplasm, site and treatments from which the transcriptional data, used in developing the GRN, was obtained. Here the *V*_c,max_ and *J*_max_ obtained in ambient [CO_2_] was used to calibrated the metabolic model, which on integration with the other models was then used to attempt to predict the observed changes in the *V*_c,max_ and *J*_max_ observed with growth at elevated [CO_2_]. Calibration was achieve by adjustinge amounts of Rubisco to match the *V*_c,max_ described for soybean grown in ambient [CO_2_]. All other protein amounts in the metabolic model, were elevated maintaining the proportion used in (Zhu et al., 2013), until a *J*_max_ was achieved that matched that observed by (Bernacchi et al., 2005). This required multiplying each by 1.2 over those used in (Zhu et al., 2013) (Supplemental table 3). The enzyme kinetic parameters of e-Photosynthesis are for 25°C. Leaf temperatures in the simulations used here slightly exceeded this. To deal with these slight, but variable, increases in temperature parameters were adjusted to the actual leaf temperature (*T*_i_) using a Q_10_ function, as described previously (Woodrow and Berry, 1988).

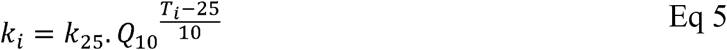

The predicted enzyme protein concentration changes as a percentage of that in ambient [CO_2_] were assumed in direct proportion to the changes of enzyme activities (*V*_*max*_) in the metabolic model. For example, if the predicted sedoheptulose-1:5 bisphosphatase (SBPase) protein concentration was predicted to increase by 3% under elevated [CO_2_], then SBPase activity (*V*_*max_Rubisco*_) was also increased by 3% in metabolic model.

#### Sensitivity analysis of each step in the metabolic model

Metabolic control analysis defines the quantitative link between the flux through a pathway and the activity of an enzyme in terms of the flux control coefficient (Fell, 1998). The maximum activity of each enzyme (*V*_*maxi*_) was both increased and decreased by 1% individually in the model to calculate the new photosynthesis rate (*A*^*+*^ and *A*^*-*^) for the two CO_2_ concentrations (350 ppm vs 550 ppm) to identify the enzymes that most influence the net photosynthetic rate. The flux control coefficient (CC) of each enzyme was calculated as:

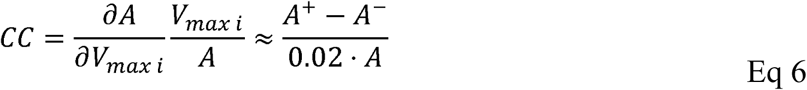

Where *A* is the original net leaf CO_2_ uptake rate, before the simulated change in activity of enzyme i (*õV*_max i_).

### Leaf level photosynthesis model

At the leaf level, the metabolic model was integrated with leaf level models of stomatal physiology, and energy balance based on the method of (Nikolov et al., 1995). Here, stomatal conductance is a function of predicted net leaf CO_2_ uptake rate, humidity, and [CO_2_] after (Collatz et al., 1991). Leaf energy balance takes account of intercepted short and long wave radiation, radiative energy loss from the leaf, convection and latent heat loss in transpiration. However, these models are inter-dependent. For example, CO_2_ uptake rate affects stomatal conductance, stomatal conductance affects leaf temperature and leaf temperature affects CO_2_ uptake rate. Solving these circular connections is achieved iteratively. Iteration continues until change to obtain a numerical solution of stomata conductance, leaf temperature, boundary-layer conductance and photosynthesis rate until changes in all four are <0.1% between iterations. This model is also implemented in MATLAB. Equations and parameters are listed in supplemental information (Supplemental table 4).

## RESULTS

### A multiscale model of soybean can mimic photosynthetic acclimation observed in FACE experiments

The integrated model predicted new steady-state enzyme concentrations for selected reactions belonging to the dark reactions and starch synthesis in the e-Photosynthesis model in response to growth of soybean leaves under elevated [CO_2_]. The ratio of predicted steady-state enzyme concentrations in elevated to ambient CO_2_ is used as one of the inputs in the e-Photosynthesis model (Supplemental Table 5). Though the magnitude of the change is small, the predicted CO_2_ response is consistent with the altered photosynthetic phenotype, as shown by the improved fit to the measured response of *A*_*sat*_ to leaf intercellular [CO_2_] and the values of *V*_c,max_ and *J*_max_ calculated from this response (Bernacchi et al., 2005) (Figure 1). Including the gene expression data, the *V*_*cmax*_ of the predicted CO_2_ response curve decreases from 115 to 109 μmol m^−2^ s^−1^, and *J*_*max*_ of the predicted curve increase from 149 to 153 μmol m^−2^ s^−1^. Simulations reveal no significant change in leaf temperature, transpiration and stomatal conductance (Supplementary Figure 1). Importantly, the fully integrated model was able to simulate the change in photosynthetic rate due to the acclimation response observed in soybean plants grown under elevated [CO_2_] (Figure 1). The predicted *A*_sat_ in 550 ppm [CO_2_] increased by 17.7% compared to ambient. Predicted metabolite concentrations also changed dynamically with elevation of [CO_2_], for example RubP decreased, PGA, T3P and SBP increased, as would be expected from an increased flux into these pools with accelerated carboxylation and decreased oxygenation at Rubisco (Figure 3).

**Figure 3.**
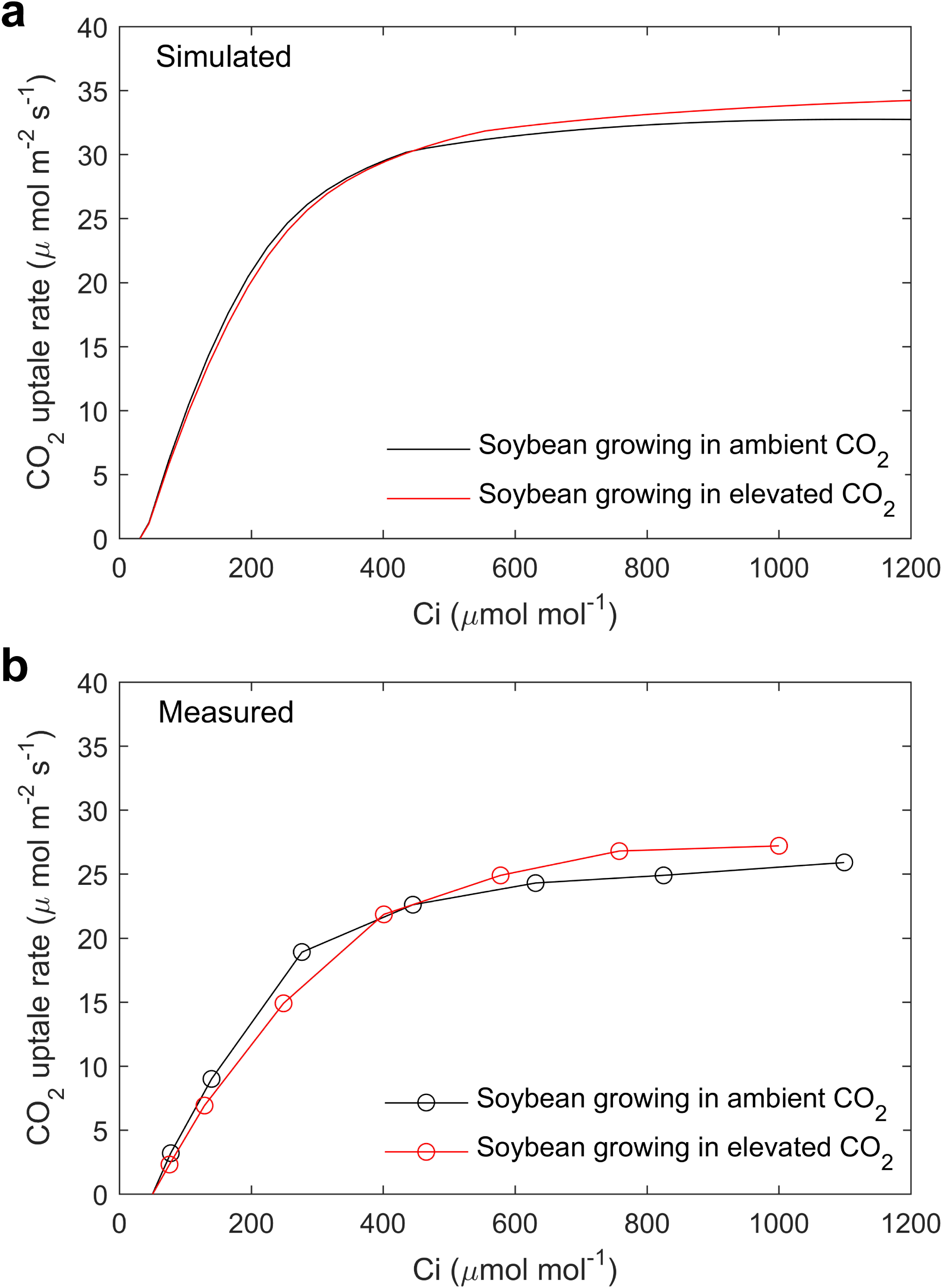
Metabolic model predicted metabolite concentrations without and with gene expression data (GE). PAR is 1200 µmol m^−2^ s^−1^.

### *in silico* perturbations reveal potential mechanisms for the transcriptional regulation of photosynthetic acclimation

Genes with a functional role in the same biological pathway are often co-expressed and sometimes co-regulated. The identification of common cis-regulatory elements in the promoters of tightly co-expressed genes is a good proxy for co-regulation (Allocco et al., 2004). The corresponding transcription factors that bind to these CREs are promising targets for the manipulation of whole pathway expression. The re-engineering of photosynthesis is needed to increase crop productivity in response to climate change (Zhu et al., 2010), such as overcoming the limitations caused by photosynthetic acclimation at elevated atmospheric [CO_2_]. With a fully integrated model of photosynthesis it was then possible to simulate the field-level photosynthetic response to genetic perturbations under both ambient (380 ppm) and elevated (550 ppm) [CO_2_]. The ideotype for elevated [CO_2_] is one in which Rubisco content is substantially decreased and controlling components of the apparatus for RubP regeneration substantially increased. This problem was approached by revisiting the individual models to identify gene targets that significantly contribute to deliver this metabolic ideotype.

#### Sensitivity of the system to individual steps within photosynthetic carbon metabolism

To determine which steps in the system exert the strongest control on *A*_sat_ in both ambient and elevated [CO_2_], a sensitivity analysis was performed by varying each parameter +/− 1% (Table 1). Control coefficients (CCs) are calculated as the ratio of change in the amount of one enzyme divided by change in *A*_sat_. If a 1% change in enzyme x results in a 1% change in *A*_sat_ CC = 1, the maximum possible, while if the change in *A*_sat_ is zero, then CC = 0, meaning that no control is exerted by that enzyme. The sum of all control coefficients should approximate to 1. Rubisco has the highest CC of all enzymes in ambient [CO_2_], while SBPase has the highest control coefficient in elevated CO_2_ (Table 1). Nine enzymes with the highest CCs overall (>0.01) were further evaluated for transcriptional regulation.

**Table 1.** The flux control coefficient of photosynthetic enzymes in ambient and elevated CO_2_.

| Enzyme | Value<br>($\mu\text{mol m}^{-2} \text{s}^{-1}$) | Meta<br>(AmbCO<sub>2</sub>) | Meta<br>(EleCO<sub>2</sub>) | Meta+GE<br>(EleCO<sub>2</sub>) | Meta+GE<br>(EleCO<sub>2</sub>)* |
| --- | --- | --- | --- | --- | --- |
| Rubisco | 120 | 0.443 | 0.121 | 0.165 | 0.108 |
| Sedoheptulose-bisphosphatase | 13.35 | 0.305 | 0.674 | 0.605 | 0.746 |
| Fructose-bisphosphate aldolase | 50.2 | 0.114 | 0.091 | 0.077 | 0.047 |
| Glucose-1-phosphate<br>adenylyltransferase | 8.01 | 0.076 | 0.058 | 0.055 | 0.035 |
| The maximum rate of ATP<br>synthesis | 6 | 0.054 | 0.047 | 0.081 | 0.051 |
| Fructose-bisphosphatases | 29.91 | 0.039 | 0.031 | 0.030 | 0.021 |
| UTP-glucose-1-phosphate<br>uridylyltransferase | 3.46 | 0.021 | 0.014 | 0.014 | 0.009 |
| Fructose-bisphosphatase (C) | 1.92 | 0.014 | 0.010 | 0.011 | 0.007 |
| Fructose-2,6-bisphosphate 2-phosphatase | 0.5 | 0.010 | 0.007 | 0.006 | 0.004 |
| Transketolase | 128.57 | 0.008 | 0.003 | 0.003 | 0.001 |
| Fructose-bisphosphate aldolase<br>(C) | 3.22 | 0.007 | 0.005 | 0.005 | 0.003 |
| Glycine transaminase | 82.37 | 0.007 | 0.002 | 0.001 | 0.001 |
| (S)-2-hydroxy-acid oxidase<br>& Catalase (CAT, EC1.11.1.6) | 43.68 | 0.001 | 0.000 | 0.000 | 0.000 |
| Sucrose-phosphate synthase | 1.67 | 0.001 | 0.001 | 0.002 | 0.002 |
| Phosphoribulokinase | 446.19 | 0.001 | -0.001 | -0.001 | 0.000 |
| glycine dehydrogenase<br>(aminomethyl-transferring) | 74.84 | 0.000 | 0.000 | 0.000 | 0.000 |
| Glycerate kinase | 171.47 | -0.001 | 0.001 | -0.001 | 0.000 |
| Glyceraldehyde-3-phosphate<br>dehydrogenase (NADP+) | 166.35 | -0.001 | 0.001 | 0.001 | 0.000 |
| Glycerate dehydrogenase | 300.29 | -0.001 | 0.000 | 0.000 | 0.000 |
| Phosphoglycolate phosphatase | 1572.6 | -0.001 | 0.000 | 0.000 | 0.000 |
| Sucrose-phosphate phosphatase | 16.65 | -0.002 | 0.000 | 0.000 | 0.000 |
| Serine-glyoxylate transaminase | 99.19 | -0.004 | -0.003 | -0.002 | -0.001 |
| 6-phosphofructo-2-kinase | 3.03 | -0.009 | -0.006 | -0.006 | -0.003 |
| Fructose-bisphosphate aldolase | 50.2 | -0.015 | -0.010 | -0.008 | -0.004 |
| Phosphoglycerate kinase | 1241.24 | -0.030 | -0.023 | -0.023 | -0.016 |
| Photosynthesis rate |  | 24.876 | 29.176 | 29.047 | 29.261 |

\* Assuming the total nitrogen (protein) resource is a constant

#### A CO_2_–responsive gene regulatory network reveals co-regulated genes

A static gene regulatory network (GRN) of the nine enzymes with the highest CCs from the e-Photosynthesis model was constructed using transcriptome data from soybean grown under ambient and elevated [CO_2_] (Leakey et al., 2009). Network nodes represent the genes that encode the nine enzymes involved in photosynthesis with the highest CC. Network edges represent regulatory interactions between TFs and putative, co-expressed target genes as determined by rank correlation of expression (MR >= 0.8 and PCC >= |0.6|) and the significant presence of CREs in the promoter of target genes for a corresponding TF gene. The static GRN was used to define the regulatory interactions that contribute to the expression of each target gene, in which each gene of interest (GOI) may have more than one TF protein regulating its expression. A linear regression modeling approach (Karlebach and Shamir, 2008) was used to determine the strength of influence, or weight (w), of each TF predicted to regulate a GOI. The resulting equations containing weighted TF-target interactions that enabled dynamic simulations with the GRN, where the expression of any TF in the network could be modified (up- or down-regulated) and result in a predicted change in expression of the target GOI (Figure 4).

**Figure 4.**
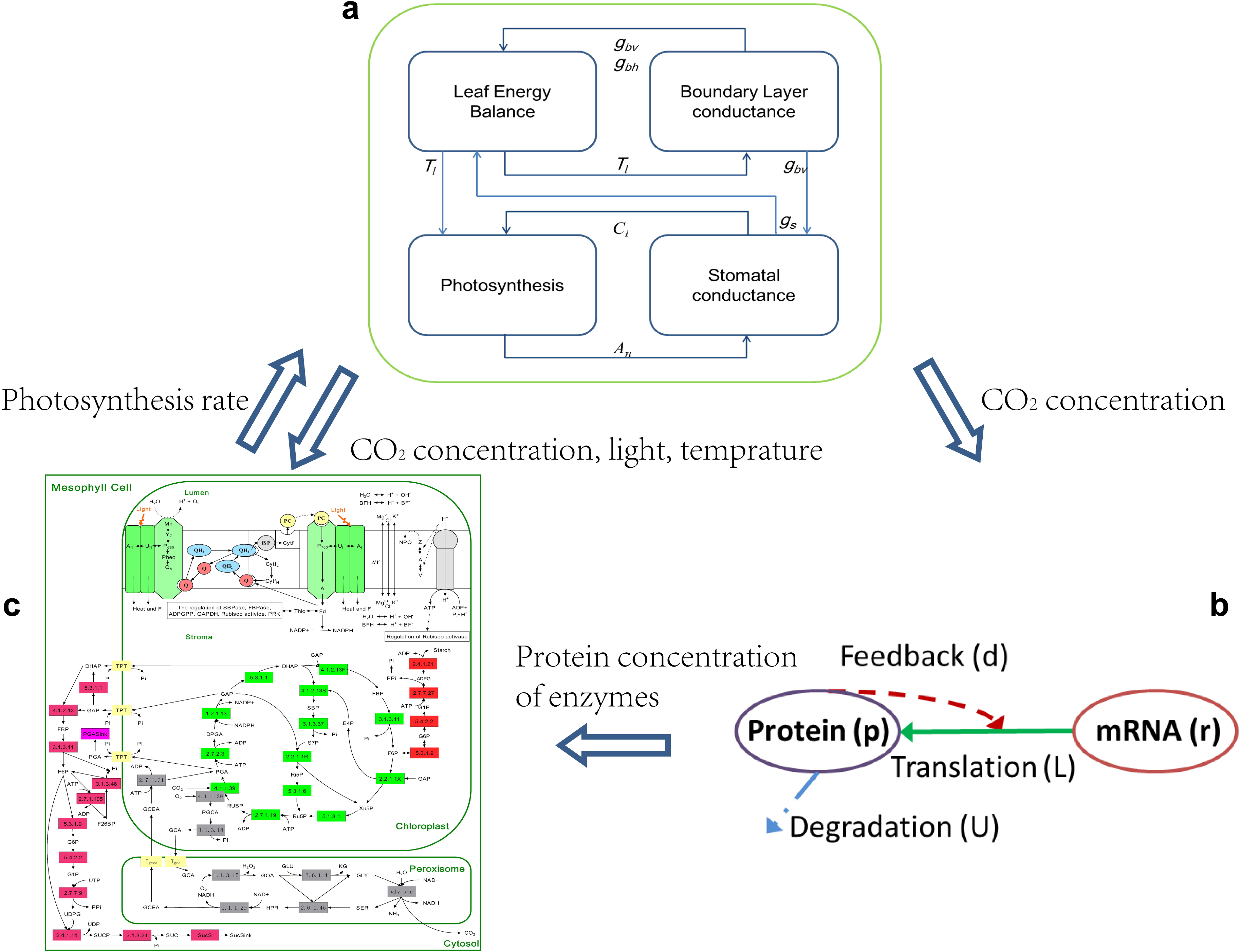
Gene expression level changes in target genes of interest as a result of *in silico* perturbation of three candidate TFs from the photosynthesis GRN. Figure shows mRNA expression levels in wild type and perturbed TF (bHLH TF knockout in a, GmWRKY71 overexpression in b, simultaneous knockout of bHLH TF and GmWRKY71 overexpression in c, GmGATA overexpression in d) conditions under ambient and elevated CO_2_.

The dynamic GRN (Figure 5) was explored to identify TFs that would simultaneously down-regulate carboxylation of Rubisco and up-regulate RubP regeneration and starch synthesis. Based on network topology, the top three candidate TFs are GmWRKY71 (Glyma.07G023300), a bHLH transcription factor (Glyma.18G115700), and GmGATA2 (Glyma.01G169400) (Supplemental Figure 2). *in silico* perturbations were performed for each candidate TF in which the TF expression was eliminated (knock-out) or overexpressed. Simulations within the dynamic GRN resulted in newly predicted expression levels of all genes targeted by the TF of interest (Figure 4).

**Figure 5.**
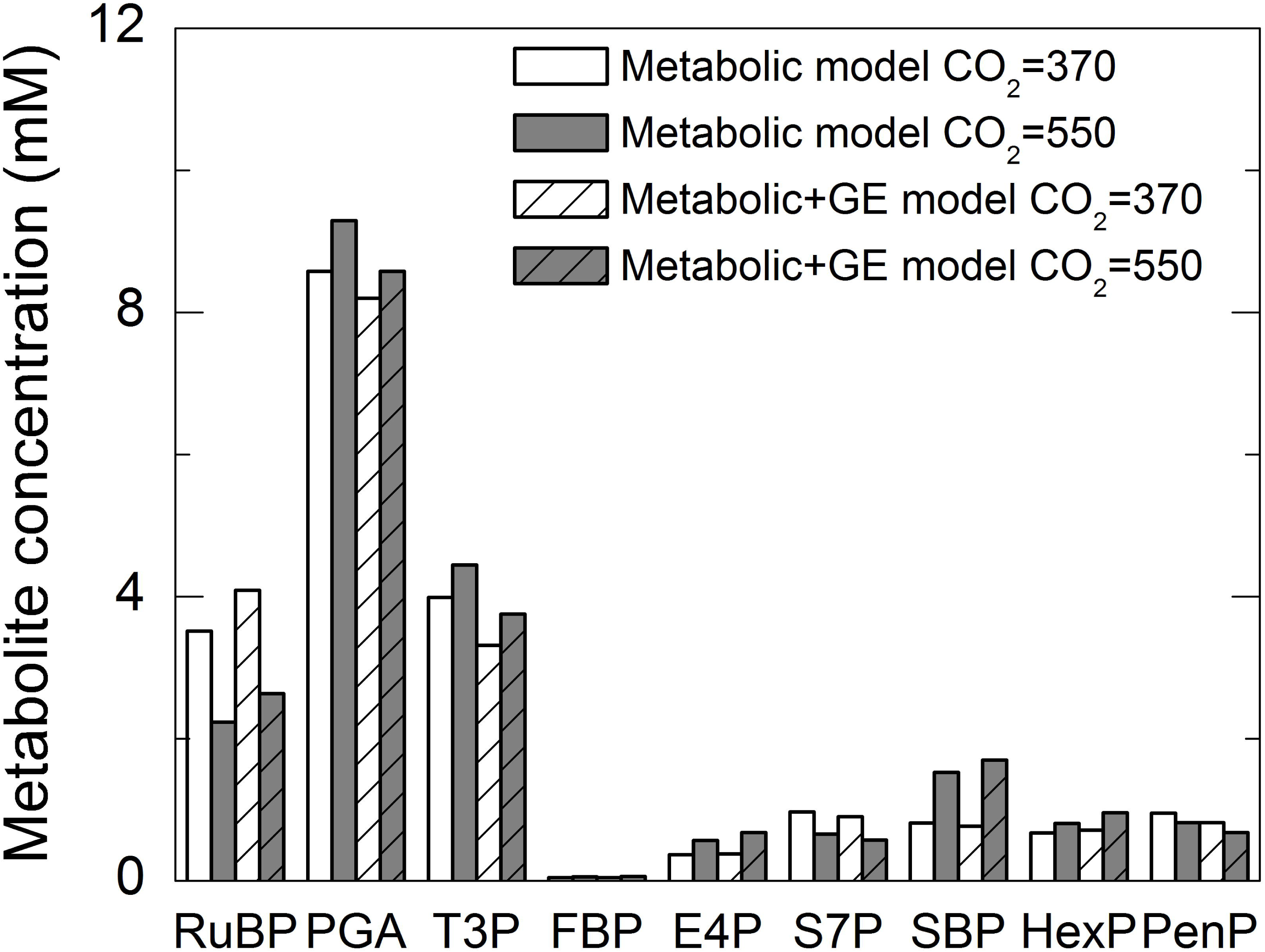
Gene regulatory network of metabolic genes having control coefficient > 0.01 based on sensitivity analysis of the metabolic model enzymes after incorporating gene expression data. Figure shows transcription factors as triangles and metabolic genes as squares. The network also shows change in mRNA expression of these genes under elevated CO_2_ concentration in leaves with blue nodes indicating downregulation and orange nodes indicating upregulation under elevated CO_2_ as compared to ambient CO_2_. Similarly, blue edges indicate predicted repression and orange edges indicate predicted activation of the metabolic gene by the TF. Thickness of the edges are based on linear model weights with more thickness indicating a heavier weight associated with the TF-target interaction.

The predicted change in mRNA expression provided input for the protein translation model that in turn predicted steady-state enzyme concentrations under elevated and ambient [CO_2_] as described in the Model Development section. The ratio of steady-state protein levels were then fed into the fully integrated photosynthesis model to obtain predicted changes in photosynthesis rate. The *in silico* over-expression of GmWRKY71 and GmGATA2, and the knockout of the bHLH TF resulted in a predicted increase in photosynthetic rate under both ambient and elevated [CO_2_]. A simultaneous *in silico* over-expression of GmWRKY71 and knock-out of bHLH TF increased the overall photosynthesis rate compared to wild type, but these failed to significantly lower Rubisco and release the resource that would be needed to elevate capacity for RubP regeneration without more resource investment (Figure 6A-C). The most promising TF candidate according to model simulations is the overexpression of GmGATA2, which results in the most dramatic change to both the down-regulation of carboxylation and up-regulation of RubP regeneration based on the simulated A/Ci curve (Figure 6D).

**Figure 6.**
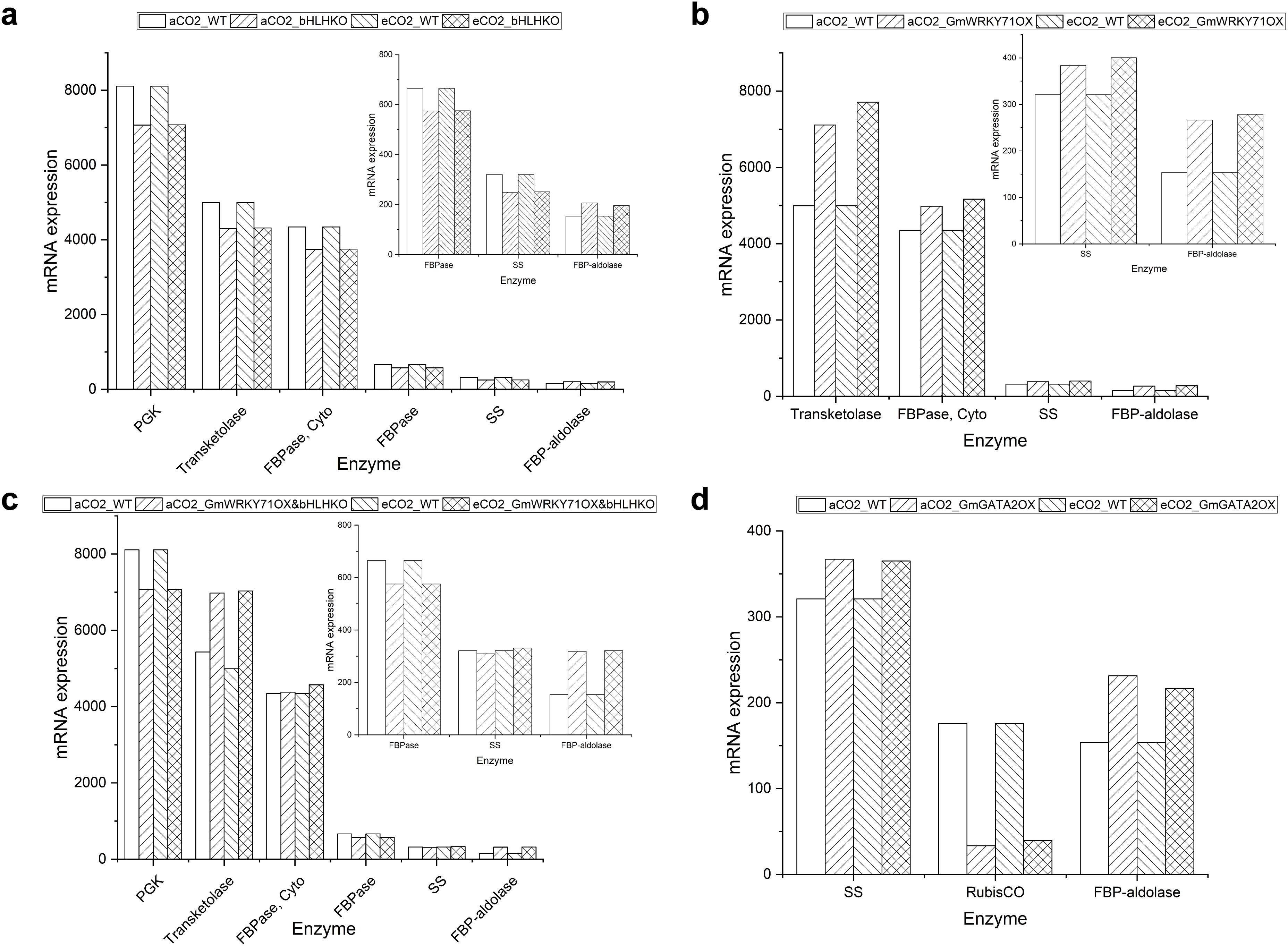
Predicted influence of transcription factor perturbations on photosynthesis rate a) bHLHB TF Knockout; b) GmWRKY71 Overexpression; c) bHLH TF Ko + GmWRKY71 Ox; d) GmGATA2 Overexpression.

## DISCUSSION

Multi-scale models have the potential to identify and add missing mechanistic details about system function and generate new hypotheses to prioritize targeted engineering efforts in plant science (Marshall-Colon et al., 2017, Millar et al., 2019). More than 4,000 mathematical plant models were published over the last decade. The majority of these models describe one biological scale or process and generalize the un-modeled spatio-temporal processes as a single output from a black box. Linking single-scale models will improve the comprehensive investigation of biological systems, resulting in explanatory models with higher prediction accuracy than models limited to one biological scale. The ability to interrogate biology at many resolutions can reveal emergent system behaviors that cannot easily be measured (Walpole et al., 2013). Multiscale models that link genes to phenotypes can accelerate the directed development of crop ideotypes. Until now metabolic modeling has predicted ideotypes with maximal photosynthetic efficiency for given environmental conditions with respect to distribution of resources between photosynthetic proteins. Achieving the ideotype then depends on identifying individual genes that might be up- or down-regulated. The multi-scale model may now be used not only to predict the metabolic ideotype, but the gene expression changes at the network level needed to achieve this. In turn, this has led, as demonstrated here, to the identification of transcription factors that can achieve the multiple gene expression changes needed by alteration of the expression of just one or two transcription factors.

In this study, we constructed a multiscale model of soybean leaf photosynthesis by integrating three models across molecular and organ-level scales using asynchronous message passing. By informing the leaf micrometeorological model with gene-level data, we were able to simulate the field-observed phenomena of photosynthetic acclimation in soybean plants grown under two different atmospheric CO_2_ concentrations. Acclimation was previously suggested to be a transcriptionally driven process that increases the capacity of respiration (Leakey et al., 2009). This though does not explain the observed decrease in in vivo Rubisco activity (Vc,max) and increase in capacity for RuBP regeneration (Jmax) by (Bernacchi et al., 2005). Existing models of photosynthesis do not provide a means to link observed transcriptional changes with metabolism and photosynthetic capacity at the leaf level. Our integrated model overcomes this and not only suggests the changes in mRNA levels and how these affect photosynthetic metabolism, but was able to predict the acclimation of photosynthetic CO_2_ assimilation that had been observed (Figure 1). Previous studies have reported proteome-level changes in response to elevated CO_2_ in soybean and other crop species (Ainsworth and Long, 2005, Bokhari et al., 2007, Taub et al., 2007, Yousuf et al., 2017, Zhao et al., 2019).

Interestingly, only 48 out of the 81 genes that encode enzymes in the e-Photosynthesis model had a statistically significant change in expression in elevated [CO_2_], and only 17 had a fold change >1.5. Since the ratio of change in enzyme concentration was proportional to the change in transcript concentration, it reveals that even a slight modification to the e-Photosynthesis model resulted in better prediction accuracy of field observations. This result represents the double-edged sword of multiscale modeling, in which fine-tuning at the molecular level can have drastic consequences at higher scales. Errors can propagate in multiscale models as information is exchanged across spatial and temporal scales. However, empirical validation of the model outputs at any scale can minimize error propagation. In our study, field data corroborated model simulations, providing confidence in the success of the modeling approach and the biologically relevant flow of information from genes to organs.

### in silico perturbations identified transcriptional regulators of photosynthesis

The successful manipulation of photosynthetic efficiency has been achieved through targeted engineering of individual enzymes (Köhler et al., 2017, Driever et al., 2017), but an alternative strategy is to simultaneously alter the expression of suites of enzymes involved in different parts of the photosynthesis pathway (Simkin et al., 2017). Understanding the transcriptional regulation of photosynthesis genes may help to fine-tune pathway expression under different environmental conditions, so avoiding the need to directly engineer change in expression of genes for each enzyme. This study uncovered three TFs, GmWRKY71, GmGATA2, and a bHLH TF, which potentially regulate the expression of genes encoding key enzymes involved in photosynthesis. While the transcriptional and post-transcriptional regulation of genes involved in photosynthesis have previously been explored (Isono et al., 1997, Fankhauser and Aubry, 2016, Saibo et al., 2008, Wang et al., 2017a, Zhang et al., 2016), this is the first report of targeted GRN analysis to identify TFs that specifically co-regulate the carboxylation of Rubisco and RubP regeneration.

Our hypothesis driven approach sought to explore how decreasing Rubisco and reallocating resources to RubP regeneration might increase *A*_*sat*_. The dynamic GRN identified high-confidence TF-target gene relationships between photosynthesis genes that have a high control coefficient and TFs tightly co-expressed with those genes. Using diverse training data, we were able to derive weights associated with each TF-target interaction, indicating which TFs exerted the greatest transcriptional control. Several TFs were found to co-regulate genes affecting Rubisco and RubP regeneration (Figure 5). However, only GmGATA2 was predicted to significantly down-regulate genes affecting Rubisco synthesis and up-regulating genes that would increase RubP regeneration and starch synthesis. The multiscale model simulations for the knockout and overexpression of GmGATA2 suggest a mechanism by which the transcriptional regulation of key photosynthetic genes can alter flux through the pathway. For example, the overexpression of GmGATA2 resulted in decreased *V*_*c_max*_ and increased *J*_*max*_, as would be required to maximize efficiency in an elevated [CO_2_] environment (Table 2; (Drake et al., 1997, Long et al., 2004)). The rewiring of metabolism, under both ambient and elevated CO_2_, produces a change in photosynthetic efficiency, which is a leaf-level phenotype. Specifically, the KO of GmGATA2 decreases overall photosynthetic capacity only in ambient [CO_2_], while overexpression results in a large decrease in carboxylation and increase in RubP regeneration according to the A/Ci curve (Figure 6D).

**Table 2.** The *V*_*cmax*_ and *J*_*max*_ of the predicted ACi curves. The *V*_*cmax*_ and *J*_*max*_ were predicted using A/Ci curve fitting utility version 2.0 (Sharkey, 2015).

| Treatment | $V_{\text{cmax}}(25^{\circ}\text{C})$ | $J_{\text{max}}(25^{\circ}\text{C})$ |
| --- | --- | --- |
| WT-Amb | 115.49 | 149.22 |
| WT-Ele | 109.17 | 152.57 |
| WT-Ele (ConN) | 112.29 | 153.58 |
| **bHLH TFOx-Amb** | 12.97 | 33.55 |
| **bHLH TFOx-Ele** | 110.66 | 90.71 |
| **bHLH TFKo-Amb** | 116.43 | 153.05 |
| **bHLH TFKo-Ele** | 114.54 | 154.51 |
| **GmWRKY71Ox-Amb** | 115.12 | 153.5 |
| **GmWRKY71Ox-Ele** | 112.08 | 154.97 |
| **GmWRKY71Ko-Amb** | 116.02 | 142.86 |
| **GmWRKY71Ko-Ele** | 113.37 | 151.82 |
| **bHLH TFKo&GmWRKY71Ox-Amb** | 118.93 | 154.76 |
| **bHLH TFKo&GmWRKY71Ox-Ele** | 114.67 | 155.78 |
| **GmGATA2Ox-Amb** | 91.28 | 157.58 |
| **GmGATA2Ox-Ele** | 89.67 | 157.59 |
| **GmGATA2Ko-Amb** | 123.28 | 135.62 |
| **GmGATA2Ko-Ele** | 119.75 | 147.49 |

ConN: Assuming the total nitrogen (protein) resource is a constant

All of the top candidate TFs from our model simulations belong to TF families that have previously been implicated in the transcriptional regulation of photosynthesis (Saibo et al., 2008). For example, bHLH TFs (Myc Family of TFs) were found to regulate aspects of C_4_ photosynthesis that are also related to genes in the ancestral C_3_ state (Borba et al., 2018). Interestingly, a number of other studies identified the GATA family of transcription factors as important regulators of photosynthesis and of carbon and nitrogen balance. The overexpression of Class B GATAs and GLKs in Arabidopsis roots improved photosynthesis by increasing root chlorophyll content (Ohnishi et al., 2018). GmGATA2 is annotated as NITRATE□INDUCIBLE, CARBON METABOLISM□INVOLVED (GNC) and is homologous to AtGATA5, and both are Class A GATAs that are associated with the light regulation of gene expression and photomorphogenesis (Zhang et al., 2015). The overexpression of the poplar PdGNC gene in Arabidopsis improved photosynthesis under low N levels by increasing the size and number of chloroplasts per cell. The photosynthetic rate in transgenics increased by 42% compared to WT lines (An et al., 2014). These studies with structurally or functionally similar GATA TFs provide support for the role of GmGATA2 in the regulation of photosynthesis. Our GRN analysis uncovered a strong positive correlation (co-expression) between GmGATA2 and FBP-aldolase and starch synthase and a strong negative correlation (anti-correlation) between GmGATA2 and the gene Rbcs that encodes the RuBisCO small sub-unit. Metabolic modeling and direct up-regulation have suggested that both FBP-aldolase and starch synthase exert strong control on RubP-regeneration (Zhu et al., 2007, Uematsu et al., 2012, Tian et al., 2018). These network predictions provide testable hypotheses for the next round of experimentation and modeling.

The multiscale modeling strategy described here represents a uni-directional flow of information from genes to physiological phenotype. However, bi-directional inputs and outputs exist between the e-Photosynthesis model and the leaf micrometeorological model, in which the metabolic model can accept environmental parameters from the organ-level model. Ideally, a truly dynamic and biologically accurate model would have a bi- or multi-directional flow of information across scales. A limitation of this model is a lack of feedback from the metabolic model to gene expression. This limitation stems from an inadequate amount of species- and condition-specific transcriptional studies. Given the availability of more expression data, it would be possible to include a model component with switch-like behavior that pulls in appropriate expression data based on environment-level inputs. Alternatively, given protein expression data it may be possible to leverage the proportional relationship between change in protein concentration and change in transcript levels to predict gene expression based on protein levels. Regardless of the method, this gap in information flow is an area of focus to improve the current model. Moreover, this multiscale model is focused on one biological process, photosynthesis. The proof-of-concept modeling approach outlined in our study provides a feasible workflow, and a base model that can be expanded on to include related pathways and processes that are still black boxes and beyond the scope of in our current model.

### Future directions

We are now poised to explore the multiscale model generated hypotheses, including the functional testing of the top TF candidate genes. Ideally, these hypotheses will be tested directly in soybean through the generation of transgenic plants that can be grown in FACE experiments (Ainsworth et al., 2008). Likewise, the model is in place for expansion to include additional metabolic pathways, but also scale to other levels. For example, the leaf micrometeorological model is already a sub-component of a canopy-level model (Drewry et al., 2010, Srinivasan et al., 2017), so an intuitive next step would be link to models that provide 3-D biophysical representations of stands of plants, as for example those developed for sugarcane agronomy (Wang et al., 2017b). This allows simulations with more realistic inputs for light capture and competition between plants. Finally, advanced visualization of multiscale model outputs will be an important next step in the simulation and analysis of *in silico* crops. Combined modeling and visualization approaches will lead to realistic simulations of ideotypes to guide selection of genetic targets for crop improvement (Marshall-Colon et al., 2017).

### Conclusion

Despite the many assumptions that have had to be made in this first linkage of gene expression networks, through protein concentrations and photosynthetic metabolism to leaf level CO_2_ exchange, it was successful in accurately predicting the observed acclimation of photosynthetic capacity in soybean when grown under elevated [CO_2_]. Most importantly it is shown to provide a numerical means to identify from many hundreds of possible transcription factors, those most likely to adapt photosynthetic efficiency to global atmospheric change. It opens the way to guiding sustainable adaptation of crop photosynthesis to a range of both current and future environments.

## ACKNOWLEDGEMENTS

Funding for this work was provided by the FFAR, awards 515760 and 602757 to AMC, SPL, and MML and from a seed grant from the Institute for Sustainability, Energy, and Environment at UIUC, and the National Center for Supercomputing Applications at UIUC. The authors would like to thank Dr. Zoi Rapti and Dr. Saurabh Sinha at the University of Illinois for their guidance in development of the translation model and dynamic GRN, respectively. We thank Dr. Venkat Srinivasan, at the University of Illinois, for valuable discussions and advice on linking leaf energy balance with CO_2_ assimilation and stomatal conductance.

## Author Contributions

AMC, SPL, YW, and KK designed the study. KK, YW, GSC performed computational analysis. MML designed the computational interface. AMC, SPL, KK, YW, GSC, and MML wrote the paper.

## SUPPLEMENTARY TABLES AND FIGURES

Supplemental figure 1. Simulated variation of assimilation (a), transpiration (b), stomatal conductance (c), and leaf temperature (d) as a function of leaf internal CO_2_ concentration under ambient CO_2_ (black) and elevated CO_2_ (red). PPFD is 1200 µmol m^−2^ s^−1^

Supplemental figure 2. Sub-networks for three transcription factors from the dynamic photosynthesis GRN chosen for *in silico* perturbation. The figure consists of bHLH TF (a), GmGATA2 (b) and GmWRKY71 (b) along with their predicted direct targets. Network nodes and interactions can be interpreted as in figure 5 of the main text.

Supplemental table 1. Least squares optimized weights for transcription factors regulating enzymes with high control coefficient after integration of protein translation model with e-photosynthesis metabolic model in the dynamic photosynthesis GRN. This table is provided separately as an excel workbook.

Supplemental table 2. Gene specific ‘d’ parameter values used in the protein translation model

Supplemental table 3. Vmax, Kcat, molecular weight and protein content used in the e-photosynthesis metabolic model

Supplemental table 4. Leaf level photosynthesis model parameters

Supplemental table 5. Steady state protein concentration ratios predicted by the protein translation model for enzymes that are part of the e-photosynthesis model

## Appendix 1 Abbreviations and units – Some variables have been added here

*A*: Net carbon assimilation (µmol m^−2^ s^−1^)

*A*_*sat*_: Light saturated *A* (µmol m^−2^ s^−1^)

*C*_*i*_: Leaf intercellular CO_2_ concentration (µmol mol^−1^)

[CO_2_]: Atmospheric CO_2_ concentration (µmol mol^−1^)

FACE: Free Air [CO_2_] Enrichment

*g*_*s*_: Stomatal conductance (mmol m^−2^ s^−1^)

*J*: Rate of electron transport (µmol m^−2^ s^−1^)

*J*_*max*_: Maximum rate of electron transport (µmol m^−2^ s^−1^)

*R*_*d*_: Mitochondrial respiration (µmol m^−2^ s^−1^)

Rubisco: Ribulose-1,5-bisphosphate carboxylase/oxygenase RubP: Ribulose-1,5-bisphosphate

SoyFACE: Soybean Free Air [CO_2_] Enrichment

*T*_*leaf*_: Leaf temperature (°C)

*V*_*c,max*_: Maximum velocity of carboxylation (µmol m^−2^ s^−1^)

