## Supplemental for "Combining gene network, metabolic, and leaf-level models show means to future-proof soybean photosynthesis under rising CO_2_"

### Supplemental tables

**Supplemental table 1:** Least squares optimized weights for transcription factors regulating enzymes with high control coefficient after integration of protein translation model with e-photosynthesis metabolic model in the dynamic photosynthesis GRN. This table is provided separately as an excel workbook.

**Supplemental table 2:** Gene specific 'd' parameter values used in the protein translation model

| Glyma_ID | Photosynthesis Pathway | Gene Symbol | D parameter value |
|---|---|---|---|
| Glyma.01G026700 | DarkRxn | GGAT | 10^-5 |
| Glyma.01G115900 | LightRxn | CAB29.1, Chloro | 10^-7 |
| Glyma.01G180800 | LightRxn | PSBP1, Chloro | 10^-6 |
| Glyma.02G064700 | LightRxn | CAB6, Chloro | 10^-7 |
| Glyma.02G080800 | LightRxn | CAB1, Chloro | 10^-6 |
| Glyma.02G094900 | LightRxn | PSBP Like-1, Chloro | 10^-6 |
| Glyma.02G249600 | LightRxn | RCA, Chloro | 10^-10 |
| Glyma.02G282500 | LightRxn | PSBP1, Chloro | 10^-7 |
| Glyma.02G309500 | LightRxn | LHCA3 | 10^-6 |
| Glyma.03G068100 | LightRxn | RCA, Chloro | 10^-6 |
| Glyma.03G114600 | LightRxn | PSBP1, Chloro | 10^-7 |
| Glyma.03G123100 | LightRxn | PPD6 | 10^-8 |
| Glyma.03G185800 | DarkRxn | T3PI | 10^-9 |
| Glyma.03G230300 | LightRxn | PSBP | 10^-6 |
| Glyma.03G253500 | LightRxn | VDE, Chloro | 10^-8 |
| Glyma.03G262300 | LightRxn | LHCA2 | 10^-9 |
| Glyma.04G014500 | LightRxn | PPD3, Chloro | 10^-7 |
| Glyma.04G015900 | DarkRxn | GAPDH | 10^-5 |
| Glyma.04G020300 | LightRxn | PETE, Chloro | 10^-6 |
| Glyma.04G051200 | DarkRxn | GDH | 10^-5 |
| Glyma.04G066000 | LightRxn | ATPSynthase | 10^-5 |
| Glyma.04G112800 | LightRxn | PSI_PSAK | 10^-8 |
| Glyma.04G167900 | LightRxn | CAB4, Chloro | 10^-11 |
| Glyma.04G209900 | LightRxn | PPD2, Chloro | 10^-7 |
| Glyma.04G215800 | LightRxn | PSAN, Chloro | 10^-7 |
| Glyma.04G241000 | LightRxn | PPD4, Chloro | 10^-8 |
| Glyma.04G242000 | LightRxn | Thioredoxin | 10^-6 |
| Glyma.04G249700 | LightRxn | PSBS, Chloro | 10^-4 |
| Glyma.05G022900 | LightRxn | PSAIII, Chloro | 10^-10 |
| Glyma.05G025300 | DarkRxn | Ru5PE | 10^-9 |
| Glyma.05G048000 | LightRxn | F-ATPase | 10^-8 |
| Glyma.05G112100 | LightRxn | FDXC2 | 10^-9 |
| Glyma.05G132600 | LightRxn | Thioredoxin O1, Mito | 10^-8 |
| Glyma.05G149400 | LightRxn | CAB | 10^-9 |

| | | | |
|---|---|---|---|
| Glyma.06G017900 | DarkRxn | GO | 10^-10 |
| Glyma.06G094300 | DarkRxn | G6PI | 10^-10 |
| Glyma.06G323700 | DarkRxn | SPS | 10^-12 |
| Glyma.07G019700 | LightRxn | PSAVI-1, Chloro | 10^-7 |
| Glyma.07G028500 | DarkRxn | Diphosphatase | 10^-8 |
| Glyma.07G142700 | DarkRxn | FBPase | 10^-7 |
| Glyma.08G044100 | DarkRxn | PGM, Cyto | 10^-8 |
| Glyma.08G074000 | LightRxn | CAB | 10^-9 |
| Glyma.08G087100 | LightRxn | Thioredoxin O1, Mito | 10^-8 |
| Glyma.08G088000 | LightRxn | TRX | 10^-7 |
| Glyma.08G097300 | DarkRxn | GO | 10^-8 |
| Glyma.08G126800 | LightRxn | FDX1, Chloro | 10^-7 |
| Glyma.08G165400 | DarkRxn | PGK | 10^-6 |
| Glyma.08G165500 | DarkRxn | PGK | 10^-8 |
| Glyma.08G173700 | LightRxn | PSBR, Chloro | 10^-7 |
| Glyma.08G302600 | DarkRxn | SGAT | 10^-8 |
| Glyma.09G087700 | LightRxn | PSAK, Chloro | 10^-9 |
| Glyma.09G154700 | LightRxn | CAB26, Chloro | 10^-8 |
| Glyma.09G250800 | LightRxn | PSAXI, Chloro | 10^-8 |
| Glyma.09G255200 | DarkRxn | HPR | 10^-8 |
| Glyma.10G042000 | LightRxn | PSAIV-A, Chloro | 10^-9 |
| Glyma.10G086600 | DarkRxn | SPP | 10^-11 |
| Glyma.10G161600 | LightRxn | TRP26 | 10^-7 |
| Glyma.10G210700 | DarkRxn | PGM | 10^-10 |
| Glyma.10G249000 | LightRxn | PSAII-1, Chloro | 10^-8 |
| Glyma.10G268500 | DarkRxn | Aldolase | 10^-9 |
| Glyma.10G293500 | DarkRxn | Transketolase | 10^-6 |
| Glyma.11G100800 | LightRxn | HCF136, Chloro | 10^-10 |
| Glyma.11G169700 | DarkRxn | PFK | 10^-10 |
| Glyma.11G226900 | DarkRxn | SBPase | 10^-7 |
| Glyma.12G199400 | LightRxn | PET | 10^-7 |
| Glyma.12G202500 | LightRxn | PSB27-1, Chloro | 10^-7 |
| Glyma.12G215100 | LightRxn | PSBQ Like - 2, Chloro | 10^-8 |
| Glyma.13G028200 | LightRxn | psbA | 10^-7 |
| Glyma.13G204800 | LightRxn | APTC1 | 10^-8 |
| Glyma.13G222300 | DarkRxn | GHMT | 10^-6 |
| Glyma.14G067000 | LightRxn | RCA, Chloro | 10^-9 |
| Glyma.15G012500 | DarkRxn | GLK | 10^-7 |
| Glyma.15G114600 | LightRxn | Cytochrome B | 10^-7 |
| Glyma.15G136200 | DarkRxn | Ri5PI | 10^-7 |
| Glyma.16G168000 | DarkRxn | FBPase | 10^-7 |
| Glyma.17G015600 | DarkRxn | AGPase | 10^-7 |
| Glyma.19G017200 | DarkRxn | G6PI, Cyto | 10^-7 |

| | | | |
|---|---|---|---|
| Glyma.19G022900 | DarkRxn | SS | 10^-7 |
| Glyma.19G046800 | DarkRxn | RuBisCO | 10^-7 |
| Glyma.19G089100 | DarkRxn | PRK | 10^-7 |
| Glyma.19G251000 | LightRxn | VDE, Chloro | 10^-8 |

**Supplemental table 3:** Vmax, Kcat, molecular weight and protein content used in the e-photosynthesis metabolic model

| EC | No. in the model | Reference | BK(1/s) | Molecular Weight (D) | Reference | Vmax (µmol m-2 s-1) | Protein content (µg m-2) |
|---|---|---|---|---|---|---|---|
| 4.1.1.39 | 1 | Zhu et al.(2007) | 16 | 588000 | Spreitzer and Salvucci (2002) | 120 | 4410000 |
| 2.7.2.3 | 2 | Zhu et al.(2007) | 540 | 45000 | Fifis and Scopes (1978), Bentahir et al (2000) | 1241.24 | 103437 |
| 1.2.1.12 | 3 | Zhu et al.(2007) | 50 | 180000 | Speranza and Ferri (1982) | 166.35 | 598860 |
| 4.1.2.13 | 5,8 | Zhu et al.(2007) | 65 | 70000 | Krueger and Sschnarrenberger (1983), Moorhead and Pplaxton (1990) | 50.2 | 54062 |
| 3.1.3.11 | 6 | Zhu et al.(2007) | 22 | 160000 | Tang et al. (2000), Reichert et al.(2000) | 29.91 | 217527 |
| 2.2.1.1 | 7,10 | Zhu et al.(2007) | 69 | 160000 | Nilsson et al (1998) Teige et al. (1998) | 128.58 | 298157 |
| 3.1.3.37 | 9 | Zhu et al.(2007) | 81 | 66000 | Cadet et al. (1988), Cadet and Meunier (1987) | 13.35 | 10878 |
| 2.7.1.19 | 13 | Zhu et al.(2007) | 216 | 96000 | Surek et al. (1985), Porter et al. (1986) | 446.19 | 198307 |
| 2.7.7.27 | 23 | Zhu et al.(2007) | 546 | 210000 | Kleczkowski et al. (1993), Li and Preiss(1992) | 8.01 | 3081 |
| 3.1.3.18 | 112 | Zhu et al.(2007) | 292 | 100000 | Kim et al. (2004), Kerr | 1572.6 | 538562 |

| | | | | | | | |
|---|---|---|---|---|---|---|---|
| | | | | | and Gear (1974) | | |
| 2.7.1.31 | 113 | Zhu et al.(2007) | 200 | 47000 | Kleczkowski et al. (1985), Kleczkowski and Randall (1988) | 171.47 | 40295 |
| 1.1.1.79 | 121 | Zhu et al.(2007) | 437 | 300000 | Kleczkowski et al. (1986), Zelitch(1955) | 43.68 | 29986 |
| 2.6.1.45 | 122 | Zhu et al.(2007) | 97 | 105000 | Ireland and Joy (1983), Paszkowski and Niedzielsa (1990) | 99.19 | 107371 |
| 1.1.1.29 | 123 | Zhu et al.(2007) | 1629 | 90000 | Julliard and Breton-Gilet (1997), Izumi et al.(1990) | 300.29 | 16591 |
| 2.6.1.44 | 124 | Zhu et al.(2007) | 54 | 90000 | Paszkowski and Niedzielska (1989) | 82.37 | 137283 |
| 1.4.4.2 | 131 | Zhu et al.(2007) | 18 | 270000 | Hiraga and Kikuchi (1980), Kochi and Kikuchi (1974) | 74.84 | 1122600 |
| 4.1.2.13c | 51 | Zhu et al.(2007) | 65 | 70000 | Krueger and Sschnarrenberger (1983), Moorhead and Pplaxton (1990) | 3.22 | 3468 |
| 3.1.3.11c | 52 | Zhu et al.(2007) | 22 | 160000 | Tang et al. (2000), Reichert et al.(2000) | 1.92 | 13964 |
| 2.7.7.9 | 55 | Zhu et al.(2007) | 400 | 53000 | Gustafson and Gander (1972), Sowokinos et al (1993) | 3.46 | 458 |
| 2.4.1.14 | 56 | Zhu et al.(2007) | 640 | 480000 | Sonnewald et al. (1993) | 1.67 | 1253 |
| 3.1.3.24 | 57 | Zhu et al.(2007) | 2500 | 120000 | Echeverria and Salerno (1994), Lunn et al. (2000) | 16.65 | 799 |
| 3.1.3.46 | 58 | Zhu et al.(2007) | 1550 | 390000 | Pilkis et al. (1987), Villadsen and | 0.5 | 126 |

| | | | | | | | |
|---|---|---|---|---|---|---|---|
| | | | | | Nielsen (2001), | | |
| 5.3.1.6 | 11 | Zhu et al.(2007) | 3440 | 53000 | Jung et al. (2000) Noltmann (1972) | 167 | 2573 |
| 5.1.3.1 | 12 | Zhu et al.(2007) | 7100 | 55000 | Chen (2000),Kopriva (2000) | 167 | 1294 |
| 5.3.1.9 | 21 | Zhu et al.(2007) | 1000 | 56000 | Lin 2009, Baumann 1988 | 33 | 1848 |
| 5.4.2.2 | 22 | Zhu et al.(2007) | 398 | 66000 | Imada 2010, Popova 1998 | 33 | 5472 |
| 3.6.1.1 | 25 | Zhu et al.(2007) | 155 | 24484 | Kapyla 1995 Sancha 2007 | 33 | 5213 |
| 5.3.1.1c | 50 | Zhu et al.(2007) | 9000 | 26500 | Alvarez 1998, Kurzok 1984 | 33 | 97 |
| 5.4.2.2c | 54 | Zhu et al.(2007) | 398 | 66000 | Imada 2010, Popova 1998 | 33 | 5472 |
| 2.7.1.105c | 59 | Zhu et al.(2007) | 9240 | 66000 | Mauricio 2003, Cabrera 2008 | 33 | 236 |
| 3.6.3.14 | Vmax11 | Zhu et al.(2007) | 285 | 325000 | Iino 2009; Nelson 1976 | 200 | 228070 |

**Supplemental table 4:** Leaf level photosynthesis model parameters

| **Parameters** | **Values** | **Unit** | **Reference** |
|---|---|---|---|
| Ball-Berry slope | 10.6 | \ | Drewry et al., 2010 |
| Ball-Berry intercept | 0.008 | mol m−2 s−1 | Drewry et al., 2010 |
| Leaf forced convection parameter | 4.3*10^(-3) | \ | Drewry et al., 2010 |
| Leaf free convection parameter | 1.6*10^(-3) | \ | Drewry et al., 2010 |
| Leaf width | 0.06 | m | Drewry et al., 2010 |
| Leaf absorptivity to PAR | 0.85 | \ | Von Caemmere 2000 |
| Rd_25 | 0.8 | µmol m-2 s-1 | Drewry et al., 2010 |
| **Environmental condition** | | | |
| Relative humidity | 0.6 | \ | |
| Wind speed | 5 | m s-1 | |
| Temperature | 25 | ºC | |
| Atmospheric pressure | 101325 | Pa | |

**Supplemental table 5:** Steady state protein concentration ratios predicted by the protein translation model for enzymes that are part of the e-photosynthesis model

| Glyma_ID | Photosynthesis Pathway | Gene Symbol | Protein Ratio |
|---|---|---|---|
| Glyma.01G026700 | DarkRxn | GGAT | 0.95111 |
| Glyma.01G115900 | LightRxn | CAB29.1, Chloro | 0.82248 |
| Glyma.01G180800 | LightRxn | PSBP1, Chloro | 0.94217 |
| Glyma.02G064700 | LightRxn | CAB6, Chloro | 0.84210 |
| Glyma.02G080800 | LightRxn | CAB1, Chloro | 0.78244 |
| Glyma.02G094900 | LightRxn | PSBP Like-1, Chloro | 0.89724 |
| Glyma.02G249600 | LightRxn | RCA, Chloro | 0.86544 |
| Glyma.02G282500 | LightRxn | PSBP1, Chloro | 0.79690 |
| Glyma.02G309500 | LightRxn | LHCA3 | 0.95426 |
| Glyma.03G068100 | LightRxn | RCA, Chloro | 0.98092 |
| Glyma.03G114600 | LightRxn | PSBP1, Chloro | 0.91823 |
| Glyma.03G123100 | LightRxn | PPD6 | 0.81338 |
| Glyma.03G185800 | DarkRxn | T3PI | 0.89471 |
| Glyma.03G230300 | LightRxn | PSBP | 0.93358 |
| Glyma.03G253500 | LightRxn | VDE, Chloro | 0.97112 |
| Glyma.03G262300 | LightRxn | LHCA2 | 0.73411 |
| Glyma.04G014500 | LightRxn | PPD3, Chloro | 0.81047 |
| Glyma.04G015900 | DarkRxn | GAPDH | 0.88690 |
| Glyma.04G020300 | LightRxn | PETE, Chloro | 0.99252 |
| Glyma.04G051200 | DarkRxn | GDH | 0.93290 |
| Glyma.04G066000 | LightRxn | ATPSynthase | 0.95148 |
| Glyma.04G112800 | LightRxn | PSI_PSAK | 0.91634 |
| Glyma.04G167900 | LightRxn | CAB4, Chloro | 0.73244 |
| Glyma.04G209900 | LightRxn | PPD2, Chloro | 0.86855 |
| Glyma.04G215800 | LightRxn | PSAN, Chloro | 0.84596 |
| Glyma.04G241000 | LightRxn | PPD4, Chloro | 0.80210 |
| Glyma.04G242000 | LightRxn | Thioredoxin | 0.97144 |
| Glyma.04G249700 | LightRxn | PSBS, Chloro | 1.00117 |
| Glyma.05G022900 | LightRxn | PSAIII, Chloro | 0.79395 |
| Glyma.05G025300 | DarkRxn | Ru5PE | 0.79528 |
| Glyma.05G048000 | LightRxn | F-ATPase | 0.90857 |
| Glyma.05G112100 | LightRxn | FDXC2 | 0.82726 |
| Glyma.05G132600 | LightRxn | Thioredoxin O1, Mito | 0.91362 |
| Glyma.05G149400 | LightRxn | CAB | 0.76962 |
| Glyma.06G017900 | DarkRxn | GO | 0.96477 |
| Glyma.06G094300 | DarkRxn | G6PI | 1.11912 |

| | | | |
|---|---|---|---|
| Glyma.06G323700 | DarkRxn | SPS | 1.31563 |
| Glyma.07G019700 | LightRxn | PSAVI-1, Chloro | 0.92325 |
| Glyma.07G028500 | DarkRxn | Diphosphatase | 1.15672 |
| Glyma.07G142700 | DarkRxn | FBPase | 0.91422 |
| Glyma.08G044100 | DarkRxn | PGM, Cyto | 1.13716 |
| Glyma.08G074000 | LightRxn | CAB | 0.61408 |
| Glyma.08G087100 | LightRxn | Thioredoxin O1, Mito | 0.71057 |
| Glyma.08G088000 | LightRxn | TRX | 0.94224 |
| Glyma.08G097300 | DarkRxn | GO | 1.15304 |
| Glyma.08G126800 | LightRxn | FDX1, Chloro | 0.97462 |
| Glyma.08G165400 | DarkRxn | PGK | 1.04209 |
| Glyma.08G165500 | DarkRxn | PGK | 0.96663 |
| Glyma.08G173700 | LightRxn | PSBR, Chloro | 0.98646 |
| Glyma.08G302600 | DarkRxn | SGAT | 0.96975 |
| Glyma.09G087700 | LightRxn | PSAK, Chloro | 0.88116 |
| Glyma.09G154700 | LightRxn | CAB26, Chloro | 0.86240 |
| Glyma.09G250800 | LightRxn | PSAXI, Chloro | 0.81260 |
| Glyma.09G255200 | DarkRxn | HPR | 1.03682 |
| Glyma.10G042000 | LightRxn | PSAIV-A, Chloro | 0.95802 |
| Glyma.10G086600 | DarkRxn | SPP | 1.17485 |
| Glyma.10G161600 | LightRxn | TRP26 | 0.95909 |
| Glyma.10G210700 | DarkRxn | PGM | 0.82748 |
| Glyma.10G249000 | LightRxn | PSAII-1, Chloro | 0.95181 |
| Glyma.10G268500 | DarkRxn | Aldolase | 1.14401 |
| Glyma.10G293500 | DarkRxn | Transketolase | 0.95906 |
| Glyma.11G100800 | LightRxn | HCF136, Chloro | 0.90122 |
| Glyma.11G169700 | DarkRxn | PFK | 0.95309 |
| Glyma.11G226900 | DarkRxn | SBPase | 0.98134 |
| Glyma.12G199400 | LightRxn | PET | 0.96645 |
| Glyma.12G202500 | LightRxn | PSB27-1, Chloro | 0.72093 |
| Glyma.12G215100 | LightRxn | PSBQ Like - 2, Chloro | 0.76407 |
| Glyma.13G028200 | LightRxn | psbA | 0.90280 |
| Glyma.13G204800 | LightRxn | APTC1 | 0.89175 |
| Glyma.13G222300 | DarkRxn | GHMT | 1.11652 |
| Glyma.14G067000 | LightRxn | RCA, Chloro | 0.89957 |
| Glyma.15G012500 | DarkRxn | GLK | 0.84189 |
| Glyma.15G114600 | LightRxn | Cytochrome B | 0.89875 |
| Glyma.15G136200 | DarkRxn | Ri5PI | 0.91715 |
| Glyma.16G168000 | DarkRxn | FBPase | 0.92231 |
| Glyma.17G015600 | DarkRxn | AGPase | 0.98852 |
| Glyma.19G017200 | DarkRxn | G6PI, Cyto | 0.74968 |
| Glyma.19G022900 | DarkRxn | SS | 0.83002 |

| | | | |
|---|---|---|---|
| Glyma.19G046800 | DarkRxn | RuBisCO | 0.94400 |
| Glyma.19G089100 | DarkRxn | PRK | 1.14634 |
| Glyma.19G251000 | LightRxn | VDE, Chloro | 0.76740 |

#### Supplemental figures

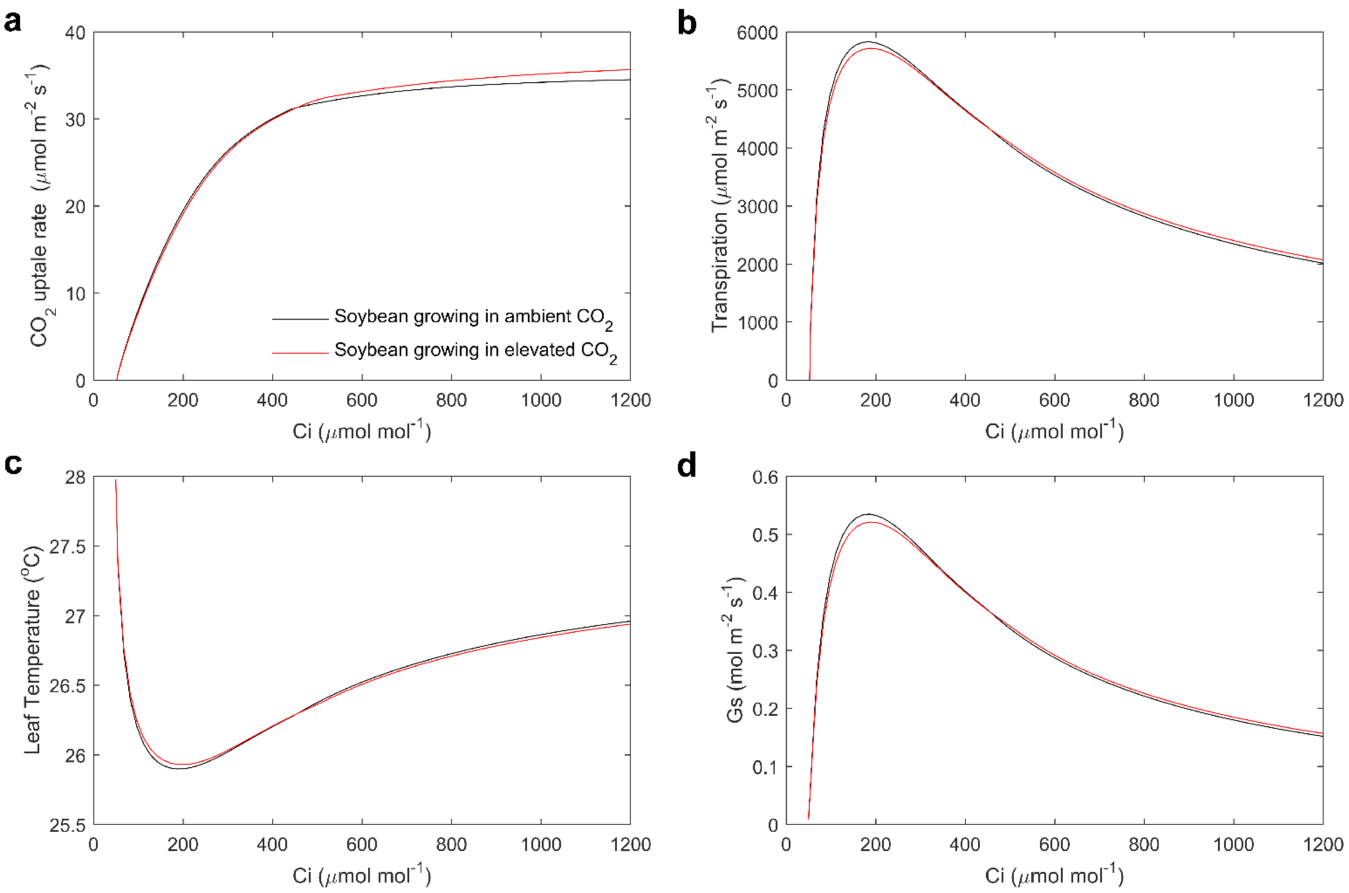


**Supplemental figure 1.** Simulated variation of assimilation (a), transpiration (b), stomatal conductance (c), and leaf temperature (d) as a function of leaf internal $CO_2$ concentration under ambient $CO_2$ (black) and elevated $CO_2$ (red). PPFD is 1200 μmol $m^{-2}$ $s^{-1}$

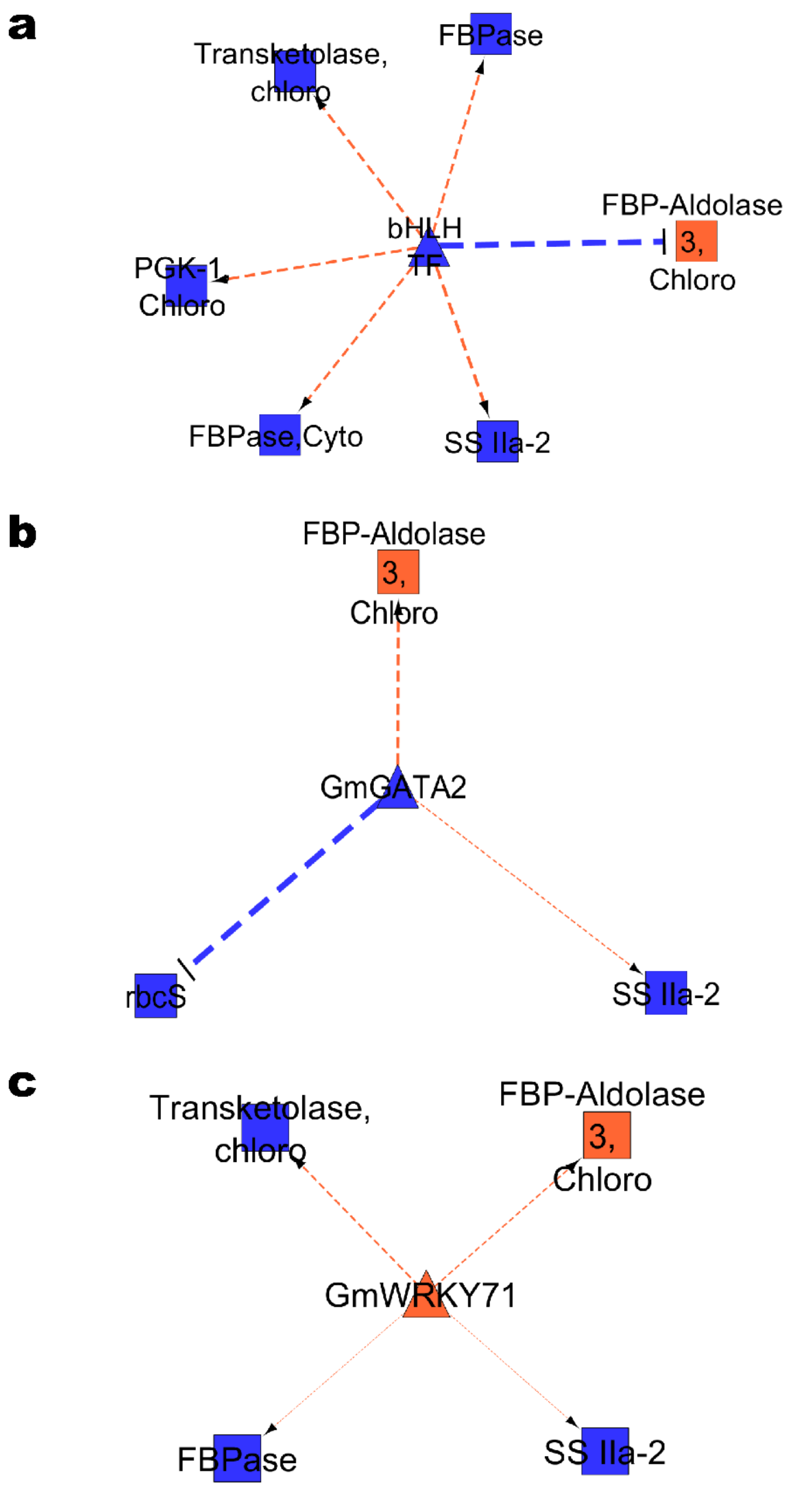


**Supplemental figure 2.** Sub-networks for three transcription factors from the dynamic photosynthesis GRN chosen for *in silico* perturbation. The figure consists of bHLH TF (a), GmGATA2 (b) and GmWRKY71 (b) along with their predicted direct targets. Network nodes and interactions can be interpreted as in figure 5 of the main text.
